# MRI Powered and Triggered Current Stimulator for Concurrent Stimulation and MRI

**DOI:** 10.1101/715805

**Authors:** Ranajay Mandal, Nishant Babaria, Jiayue Cao, Kun-Han Lu, Zhongming Liu

**Affiliations:** Weldon School of Biomedical Engineering; Purdue Institute for Integrative Neuroscience; School of Electrical and Computer Engineering, Purdue University, West Lafayette, IN, USA; MR-Link LLC, West Lafayette, IN, USA

## Abstract

Bioelectric stimulation during concurrent magnetic resonance imaging (MRI) is of interest to basic and translational studies. However, existing stimulation systems often interfere with MRI, are difficult to use or scale up, have limited efficacy, or cause safety concerns. To address these issues, we present a novel device capable of supplying current stimulation synchronized with MRI while being wirelessly powered by the MRI gradient fields. Results from testing it with phantoms and live animals in a 7 Tesla small-animal MRI system suggest that the device is able to harvest up to 72 (or 18) mW power during typical echo-planar imaging (or fast low angle shot imaging) and usable for stimulating peripheral muscle or nerve to modulate the brain or the gut, with minimal effects on MRI image quality. As a compact and standalone system, the plug-and-play device is suitable for animal research and merits further development for human applications.

## Introduction

Neurostimulation, e.g. Deep Brain Stimulation (DBS), Vagal Nerve Stimulation (VNS), Spinal Cord Stimulation (SCS), has been widely used to treat Parkinson’s disease [1], dystonia [2], [3], epilepsy [4], [5], and intractable pain syndrome [6], [7], to name a few examples. It has also been increasingly recognized that magnetic resonance imaging (MRI) can guide neuromodulation and improve its efficacy, especially if neural stimulation and imaging are performed simultaneously [8]– [12]. However, concurrent stimulation and MRI is non-trivial. A conventional stimulation device may jeopardize patient safety and degrade imaging quality [13], while MRI may interfere with the device and corrupt its function [14]– [16], due to the strong and varying magnetic fields during MRI [17]. In addition, a device often requires long cables to connect to a power source or receive external triggers, where-as such cables may perturb the magnetic fields and degrade image acquisition [18], [19].

Widely used methods for simultaneous MRI and neuromodulation involve non-MR-safe stimulators, which are placed outside the MRI room and connected to the subject using long twisted cables [20]– [23]. The longer stimulation cables sometime pick up fast gradient magnetic fields and lead to unwanted electrical stimulation [24][25]. Few studies include conditionally MR-safe stimulators placed near the MRI bore with shorter cables and RF filtering [26]– [28][29]. However, such setups are difficult to scale up, since more stimulation channels would require more cables and RF filtering circuits and amount to an increasingly bulky and complex system. The system also causes patient discomfort, prolongs preparation time, affects imaging quality, especially at a high field (7 Tesla or above) [30].

A potential solution to this problem is to use MRI to wirelessly power and drive a device. Wireless power harvesting, although common in various electrical systems [31], [32], has been rarely used to extract power from MRI. Inside an MRI system, RF fields [33] and fast switching gradient may supply energy to drive low-power electronic devices [34]. However, prior studies have only shown wireless power harvesting up to 7mw power from both RF and gradient fields [33], [34]. Apart from the powering, it is beneficial to synchronize the timing between stimulation and MRI. Wirelessly picking up magnetic-field changes can sense the timing of MRI and supply triggers to synchronize MRI with another device [35] or imaging system [36]. However, prior studies explore either wireless powering or synchronization (but not both) and use circuits and systems with large footprints, which are unsuitable for placement inside MRI bore.

To address these issues, we introduce a low-power, programmable device both powered by and synchronized with MRI. This device, herein referred to as XON, can provide biphasic electrical currents to stimulate muscle, nerve, or organ of interest during concurrent MRI, e.g. with echo-planar imaging (EPI) and Fast Low Angle Shot (FLASH) pulse sequences. As shown in Fig. 1, XON, placed inside the MRI bore, operates without any battery or cables, and offer a plug-and-play solution to simultaneous imaging and stimulation for various applications. Hereafter, we describe the system design and implementation, and present the proof-of-concept results obtained from benchtop, phantom, and in vivo animal experiments.

**Fig. 1.**
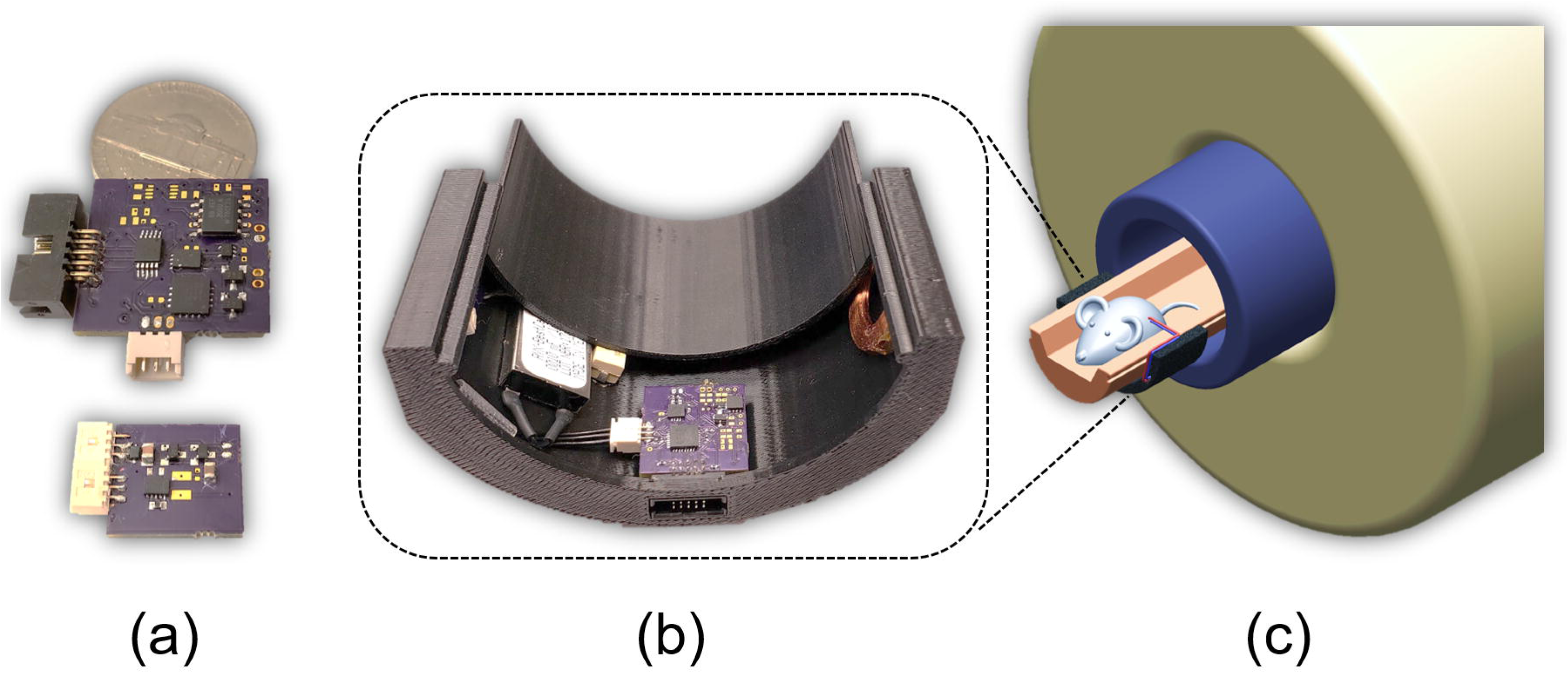
XON stimulator: (a) circuit boards. (b) 3D printed enclosure containing device elements. (c) The enclosure readily clamps onto the animal holder inside the MRI bore.

## Materials and Methods

### A. System Design

XON included (a) a programmable stimulation circuit to provide biphasic and charge-balanced current, (b) a power-harvesting circuit to scavenge wireless energy from MRI gradient fields and (c) a gradient detection circuit for synchronization of imaging and stimulation. The wireless device was able to harvest power during MRI scans, detect the timing of the MRI pulse sequence, generate electrical current, synchronized with MRI, and thus support precise stimulation during concurrent MRI. To program the device (e.g. changing the stimulation parameters), we also implemented a base station and a custom-built software to connect the device via a USB interface to a PC.

#### 1. Stimulation circuit

The stimulation circuit (Fig. 2) consisted of four major components: (a) a H-bridge to generate biphasic voltage from a single supply (b) a reference current source and sink (REF200, Texas Instruments, USA), (c) an adjustable current up-scaler and (d) isolation switches. The harvested single supply DC voltage was utilized separately for positive and negative phases of biphasic stimulation by using a H-bridge. Fig. 2 illustrates the high current (20mA) and high voltage (20V_pp_) compliance electrical stimulator’s front-end architecture. Harvested supply voltage lines (+V and Gnd) were connected to the switches S1 and S2 (ADG1436, Analog Devices Inc., USA). These switches determined the instantaneous direction of the current stimulation pulses (positive, negative, or no stimulation). The reference current generator, I_S_, employed a floating current source and Schottky diodes to produce a known current (e.g. 100 µA), which served as an input to the adjustable current up-scaler. The reference current flowed through a programmable resistor R_SET_ (AD5290, Analog Devices Inc., USA) and the load impedance. The voltage drop was buffered by an operational amplifier and passed through resistor R_O_ and the load. The resistor ratio helped up-scale the reference current from. 1mA up to 20mA. The output current flowing through the load was expressed as

**Fig. 2.**
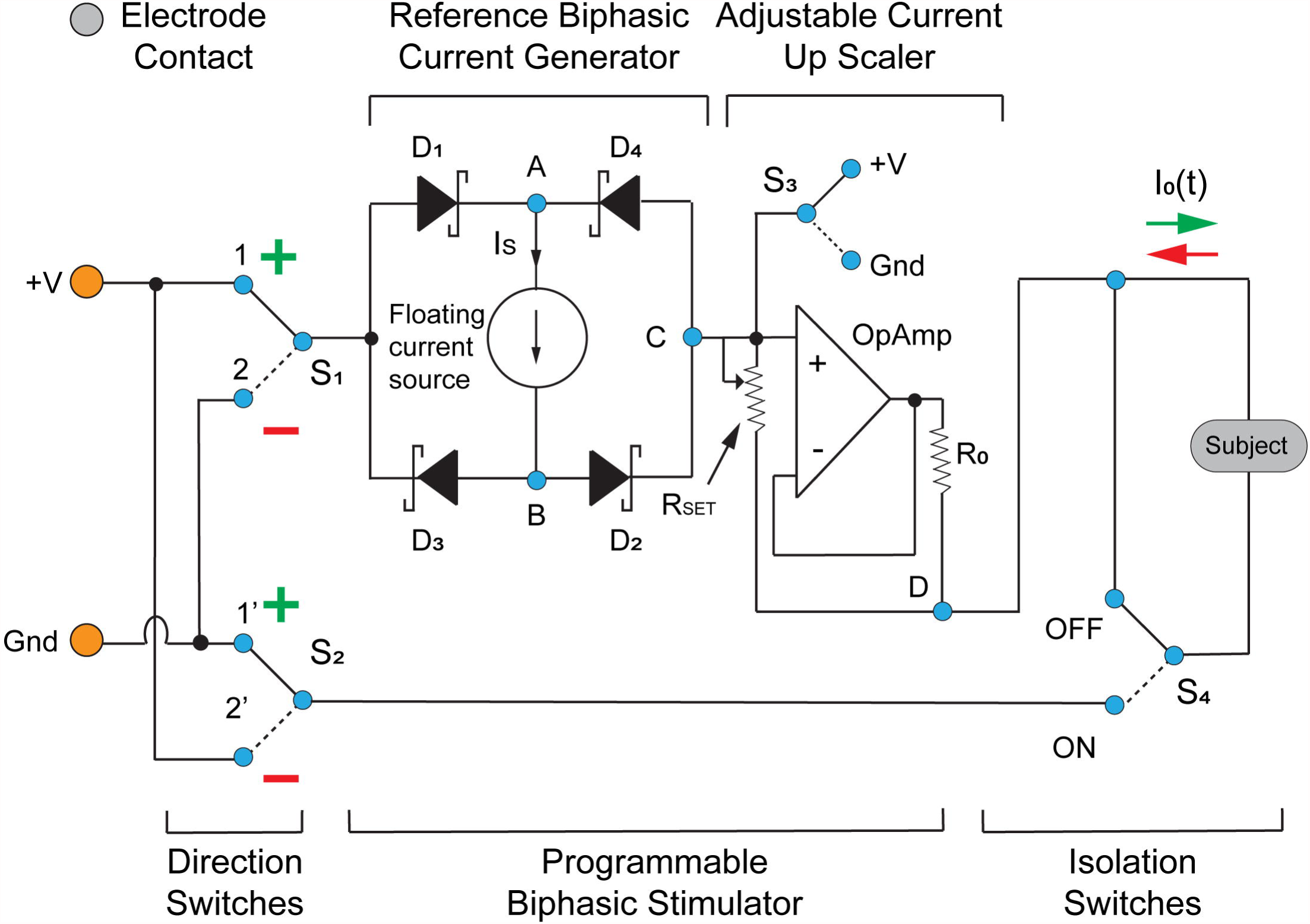
Schematic diagram for the current stimulator front end circuit.

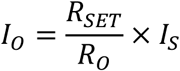

Switch S3 was utilized to determine the known voltage state at the output of Op-amp while S1 and S2 are OFF or tri-stated. The output switch S4 (ADG1419, Analog Devices Inc., USA) helped to isolate the device from the biological tissue during non-stimulation periods. The analog switches had low on-resistance (<10 Ω) and low charge-injection (~20pC) for reducing on-state voltage drop and feed-through noise.

An ultra-low-power microcontroller (MSP430F2132, Texas Instruments, USA) was used to control individual switches (S1, S2, S3 and S4) and the digital potentiometer (AD5290, Analog Devices Inc., USA). The micro-controller firmware was precisely designed to leverage the low power modes of the system. As a result, the MCU required less than 0.4 mA during stimulation operation and less than 60µA during the idle mode. The switching elements, op-Amps and digital potentiometer were chosen based on their ultra-low quiescent current. Each of these components were operated in idle modes during non-stimulation periods to ensure quiescent power consumption of the whole device during stimulation experiment was less than 1mW.

#### 2. Gradient detection circuit

The gradient detection circuit utilized a flexible PCB coil (Fig. 3b) and its tuning circuit to pick up the varying magnetic field during trapezoidal gradient pulses of fMRI. The differential signal picked up by the coil was passed through a differential passive filter and subsequently converted into a single ended signal by an instrumentation amplifier (INA826, Texas Instruments, USA). The single ended signal went through a single stage low pass filter before it was converted to binary signals (hereafter referred to as the gradient triggers) by two comparators (Fig. 3c). Each of these comparators converted the varying and steady states of the gradient magnetic field into respectively binary ‘0’ and ‘1’. The two binary signals contained information regarding the positive and negative ramping periods of the trapezoidal gradients. These gradient triggers were then fed into the microcontroller which controlled the stimulation patterns in synchrony with MRI.

**Fig. 3.**
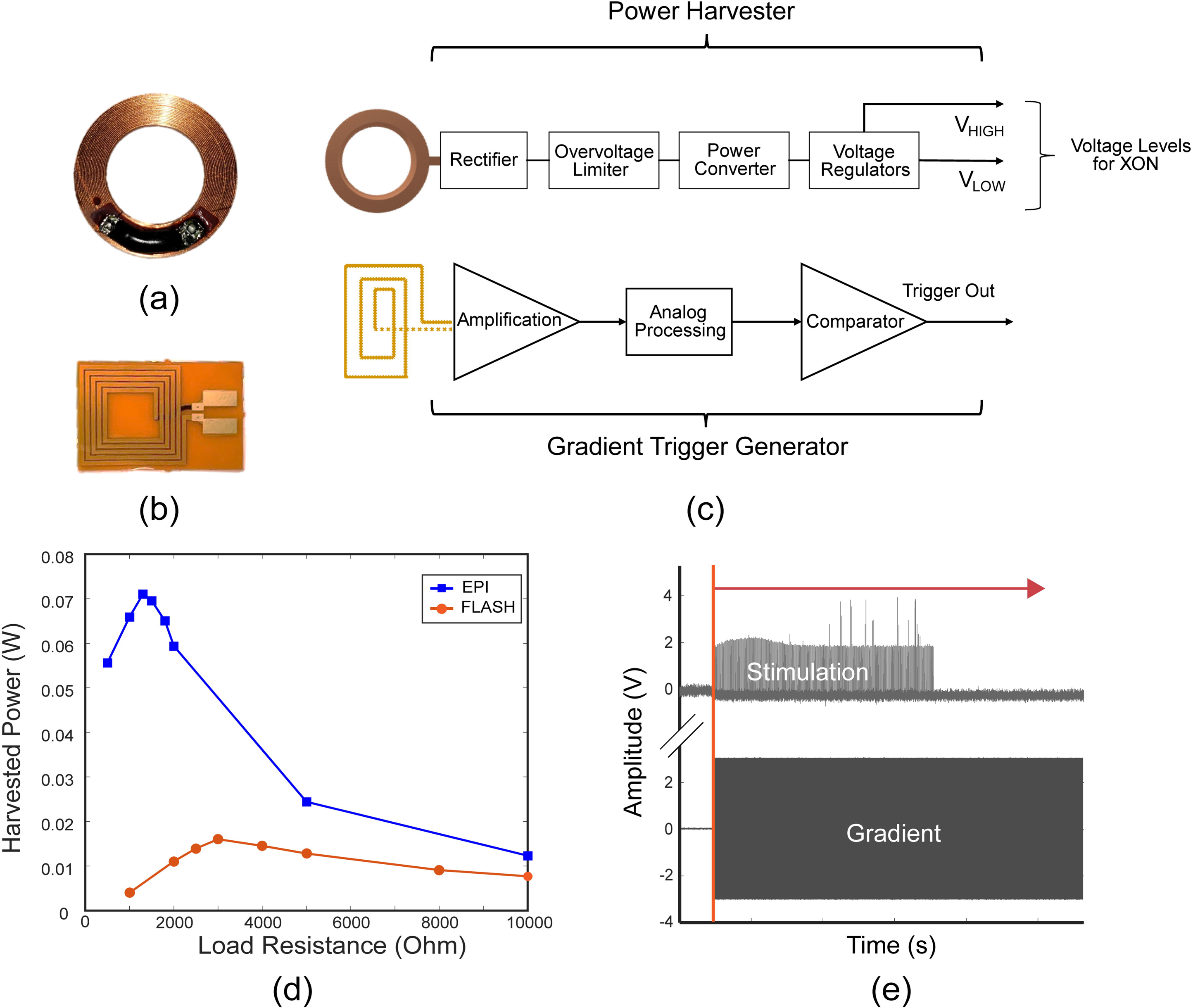
Gradient Power Harvesting and Image Trigger Generation: (a) A small machine wound coil (2cm dia.) and (b) a planar coil on a flexible PCB, was used to pick up switching gradient fields for power harvesting and imaging trigger generation. (c) Functional block diagrams of the system. (d) Wireless power from MRI gradients varied with load impedance and imaging protocols. Peak power of 72mW, harvested during EPI and 18mW during FLASH imaging. (e) Stimulation was synchronized to begin at the start of each imaging sequence.

#### 3. Power harvesting circuit

The power harvesting unit comprised of a machine wound coil (Fig. 3a) and a power management unit to generate two voltage levels for the control circuitry (+3.3V, V_LOW_) and stimulation circuitry (+11V, V_HIGH_) respectively. The machine wound coil had an inductance of 5.3uH, a series resistance of 36.5 Ohm (R_DC_) and a diameter of 2 cm. In the power management unit, the harvested AC voltage across the coil was rectified by using a full-bridge rectifier with Schottky diodes for low forward voltage drop. The DC voltage was used to charge a small ultra-capacitor (BZ05, AVX Corporation) that served as a buffer to store the harvested energy. In addition, it acted as a filter to produce a clean DC supply voltage. A Zener diode (MM3Z12V, ON Semiconductor, USA) was used to limit the voltage across the capacitor to 11V. A voltage-controlled analog switch then connected the DC voltage from the capacitor to a switching voltage regulator (TPS62177, Texas Instruments, USA), which provided regulated +3.3V supply to the microcontroller. The control voltage of the system was kept at 6V to ensure the energy stored in the capacitor was enough to deliver the required startup current to the microcontroller.

#### 4. Enclosure and assembly

Individual parts were assembled with MR-compatible cables and connectors. An enclosure (Fig.1b) was designed and 3D-printed to readily clamp on the animal holder and could be fitted easily inside the MRI bore (Fig.1c). The enclosure containing XON stimulator and coils were placed 6cm away from the imaging ROI (head or gut) to pick-up electrical energy from gradients. A ten-channel flexible ribbon cable (FRC) was used to connect the electronic circuit within the enclosure to the base station during stimulator parameter programming.

#### 5. Base station

The external base station was designed to achieve simple bi-directional communication (UART: Universal Asynchronous Receiver/Transmitter) with the onboard microcontroller of the stimulation unit. It consisted of (a) a USB to UART communication IC, (b) a power management unit, (c) a low-power microcontroller. The base station was connected to a PC via a USB cable. A custom GUI was developed in LabView (National Instruments, USA) to set up stimulation settings and parameters (e.g., frequency, amplitude, direction). Each of these parameters was sent from the PC through a 2-byte command protocol. The command was then received by the base-board microcontroller to communicate with the stimulation unit. Additionally, the base station offered four analog-to-digital conversion channels (10-byte ADC) to record auxiliary information optional to the device operation. The four channels had a combined bandwidth of 50 kHz. The digital data were transferred to the PC at 921600 bps (~1 Mbps), following the 2-byte command protocol. The power management unit converted the USB-bus voltage (5 V) into two regulated voltage outputs of 11 V and 3.3 V, which were only used for benchtop testing and disabled for experiments inside MRI.

### B. Experiments

The power-harvesting capability of the system was tested by evaluating the load power characteristics of the system during operation of two widely used MRI sequences: EPI and FLASH. The animals were scanned in a 7-tesla horizontal-bore small ani-mal MRI system (BioSpec 70/30; Bruker Instruments, Billerica, USA) equipped with a gradient insert (maximum gradient: 200mT/m; maximum slew rate: 640T/m/s), a volume transmit and receive 1H RF coil (86 mm inner-diameter) and a surface receive-only 1H RF coil (10 mm inner-diameter).

The low power stimulator was operated simultaneously during these imaging sessions to provide MRI powered and triggered stimulation. Efficacy of the wireless stimulator was analyzed through two experimental setups (a) blood oxygen level dependent (BOLD) response evoked by forepaw stimulation (with EPI) and (b) gastric motility in response to vagus nerve stimulation (VNS) (with multi-slice FLASH).

For the animal studies, two Sprague Dawley rats (male, 350-450g, Envigo RMS, Indianapolis, IN) were used for in vivo experiments. One animal was used for simultaneous fMRI and forepaw stimulation; the other animal was used for gastric MRI given VNS. All experiments were performed according to a protocol approved by the Purdue Animal Care and Use Committee and the Laboratory Animal Program.

For both experiments, animals were anesthetized with the same protocol. Each animal was initially anesthetized with 5% isoflurane for 5 minutes. After the animal was placed inside the MRI, anesthesia was maintained with continuous administration of 2–3% isoflurane mixed with oxygen. The heart and respiration rates and oxygen saturation level (SpO_2_) were monitored by using a small-animal physiological monitoring system (Kent Scientific, CT, USA). The animal’s body temperature was maintained at 37 ± 0.5°C using a water heating system.

After performing toe pinch, to assure adequate anesthesia MR-compatible needle electrodes were placed on the left forepaw of one animal. The electrodes were connected to the XON stimulator with a custom connector in preparation for BOLD fMRI and concurrent muscle stimulation.

One animal underwent neck surgery for implantation of a bipolar cuff electrode around the left cervical vagus nerve. After ad-ministering a preoperative bolus of carprofen (10mg kg^−1^, IP; Zoetis, NJ, USA) and performing a toe-pinch test, a ventral midline cervical incision was made between the mandible and sternum. The subcutaneous tissue was then dissected and retracted later-ally together with the mandibular salivary glands to reveal the trachea and the left carotid artery. Upon exposure of the left carotid artery, the left cervical vagus nerve, located lateral to and in parallel with the carotid artery above the level of the carotid bifurcation, was identified. The connective tissues surrounding the left cervical vagus nerve were carefully dissected so that a 10-15 mm portion of the cervical vagal trunk was isolated away from the carotid artery. A bipolar cuff electrode (MicroProbes, Gaithersburg, MD, USA) with a platinum-iridium wire lead was wrapped and secured on the isolated vagus nerve. The lead was externalized prior to suturing the incision site.

#### 1. Phantom experiment

In the phantom experiment, a test tube containing CuSO4 x 2H2O (1g/L) solution (Bruker Instruments, Billerica, MA) was used. A custom 3-D printed model (26×26×40 mm^3^) was placed inside the test-tube (Fig. 6f) to test whether the device caused any geometrical distortion or SNR degradation. The test tube was then positioned inside the MRI and scanned with EPI or FLASH (image sequence describer later). During imaging, the power harvesting unit was placed inside the MRI to quantify the maximum output power and a resistive load impedance was varied from 500Ω up-to 100kΩ.

#### 2. BOLD fMRI and muscle stimulation

First, we evaluated the performance of the wireless stimulator in harvesting energy from fast gradients of an EPI sequence. A 2-D single-shot gradient echo EPI sequence (TR = 1s, TE = 15ms, FA = 55°, in-plane resolution about 0.6×0.6 mm^2^, slice thickness = 1 mm) was used for imaging. Stimulation current was delivered according a block-design paradigm that consists of 30s-ON-30s-OFF blocks, programmed by the X-ON current stimulator. The stimulation was triggered to start, beginning of each imaging cycle. When the stimulator was turned ON, monophasic square pulses (pulse width: 5ms; amplitude: 1.0 mA; frequency: 10 Hz) were delivered to the animal for a total of 10 mins or 10 cycles.

#### 3. Gastric MRI and vagal nerve stimulation

To further understand the efficacy of the device, we operated it during FLASH imaging. A 2-D FLASH sequence (TR = 11.784ms, TE = 1.09ms, FA = 25°, in-plane resolution = 0.47×0.47 mm^2^, slice thickness = 1.5mm) was chosen for this experiment and a volume coil was used for recording larger abdominal area. An implanted bipolar cuff (described in earlier section) electrode was connected with the X-ON current stimulator. The stimulation current was delivered in 20s-ON-40s-OFF cycles. When the stimulator was turned ON, monophasic square pulses (pulse width: 0.5 ms; amplitude: 0.4 mA; frequency: 20 Hz) were delivered for a total of 10 cycles.

#### 4. Effects on MRI data quality

The performance of the device’s wireless power harvesting scheme on MR images were quantitatively assessed by analyzing the temporal signal-to-noise ratio (tSNR) of the fMRI data and bold activation map with forepaw stimulation. The tSNR histograms were generated in two cases: (1) where the X-ON device operated inside the MRI bore during which being wirelessly powered by the MRI and (2) when the device was placed outside of the MRI and was turned OFF. Furthermore, contrast-to-noise ratio (CNR) was computed as harvested power levels were varied. The CNR computation was repeated for FLASH sequences as well.

### C. Post-processing

The fMRI data were processed using Analysis of Functional Neuroimages (AFNI, [37]) and the FMRIB Software Library (FSL, [38]). The fMRI data were first corrected for motion and slice timing before being spatially smoothed as per one of the prior works [9], [39]. The fMRI time series were detrended by regressing out the linear trend voxel by voxel. To calculate tSNR for every voxel, the signal mean was divided by its standard deviation. The stimulation blocks were convolved with a canonical hemodynamic response function (HRF) and this model was correlated with the fMRI signal at every voxel to generate the corresponding activation map (p<0.05, uncorrected). The gastric MRI data were processed and registered using a custom-built software developed in MATLAB as described else-where [40]. The antral region of the stomach was segmented, and a contraction time series was obtained for every location along the antral axis, through a methodology described in Lu et al. [41]. The time series represented the cross-sectional area change of a particular plane. To quantify the effect of VNS on gastric motility, we extracted the contraction time series from the middle antrum.

## Results

### A. Phantom Experiment

The phantom experiment outlined a repeatable setup to characterize the X-ON device. Initially, the power harvesting unit was placed inside the custom-made enclosure and kept inside the MRI bore. The phantom test-tube was then imaged with an EPI and FLASH sequence respectively. Each of these scans were repeated several times as the load impedance across the harvested voltage rail (+11V) was varied. The maximum harvested power varied with the load impedance as shown in Fig. 3d. The peak harvested power (72mW and 18mW) during both imaging protocols were sufficient for the XON stimulator. Higher levels of power were generated during EPI sequence as compared to a FLASH sequence. This variation can be correlated with the faster and more rapid change of gradients during EPI sequence.

The stimulator front end circuit was then tested during phantom experiments as the XON device was powered through the harvesting unit. The biphasic current and voltage waveform were measured by using a resistor capacitor load impedance (Randle’s Equivalent Circuit [42]) that represents an electrode electrolyte interphase (Fig. 4c). The wireless stimulator was connected with the load impedance with connector T1 and T2. The stimulator provided load currents and voltages as shown in Fig. 4b and Fig. 4d, respectively.

The XON stimulator could be programmed with different stimulation block designs to further evaluate its performance. For an example a 30s-ON-30s-OFF bi-phasic block stimulation paradigm can be seen in Fig. 4a.

**Fig. 4.**
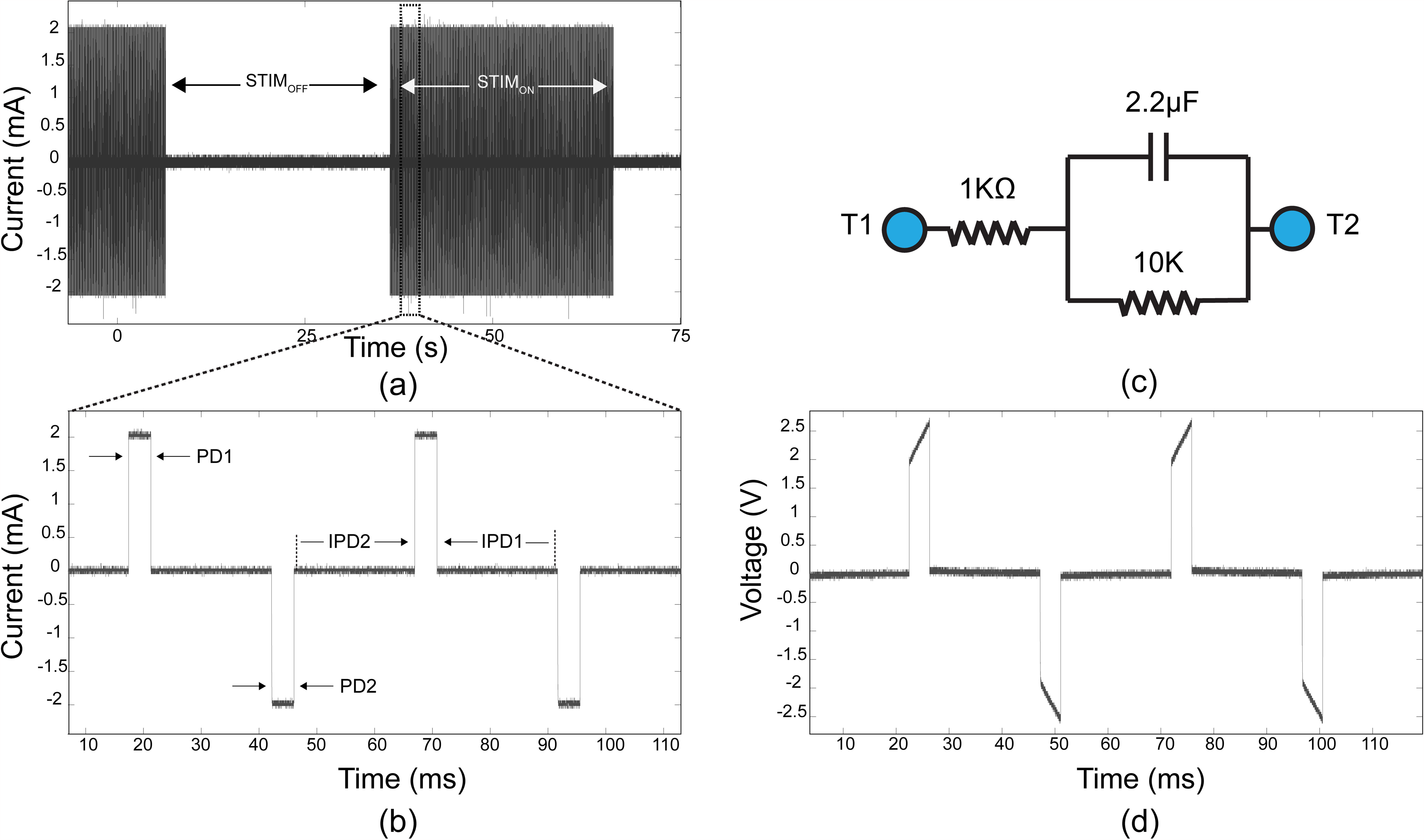
Current stimulator performance: evaluated on a Randle’s equivalent load impedance (c) of electrode-electrolyte interphase. Biphasic current pulses (b) generated through XON and resulting lead voltages (d) across T1 and T2 were measured. A 30s-ON-30s-OFF stimulation paradigm (a) was programmed into the stimulator. Individual parameters like STIMON, STIMOFF, Inter Pulse Durations (IPD1 &IPD2) and Pulse Durations (PD1 & PD2) can be tuned through the software to change the frequency, duration, polarity and amplitude of the stimulation current waveforms.

### B. BOLD fMRI and muscle stimulation

The device’s efficacy for simultaneous muscle stimulation during BOLD fMRI was evaluated in a single animal. The fast EPI gradients provided the necessary energy to sustain the device operation as the left forepaw of the rat was stimulated. The experimental paradigm can be seen in the Fig. 5a. The simultaneously acquired fMRI signal of each voxel was correlated with the stimulation block triggers and was evaluated for statistical significance (p<0.05). The correlation map clearly revealed contralateral BOLD activation (Fig. 5b) centered around the right somatosensory cortex (R-S1FL) given left forepaw stimulation. Statistically significant S1 activation was readily observed (Fig. 5b).

**Fig. 5.**
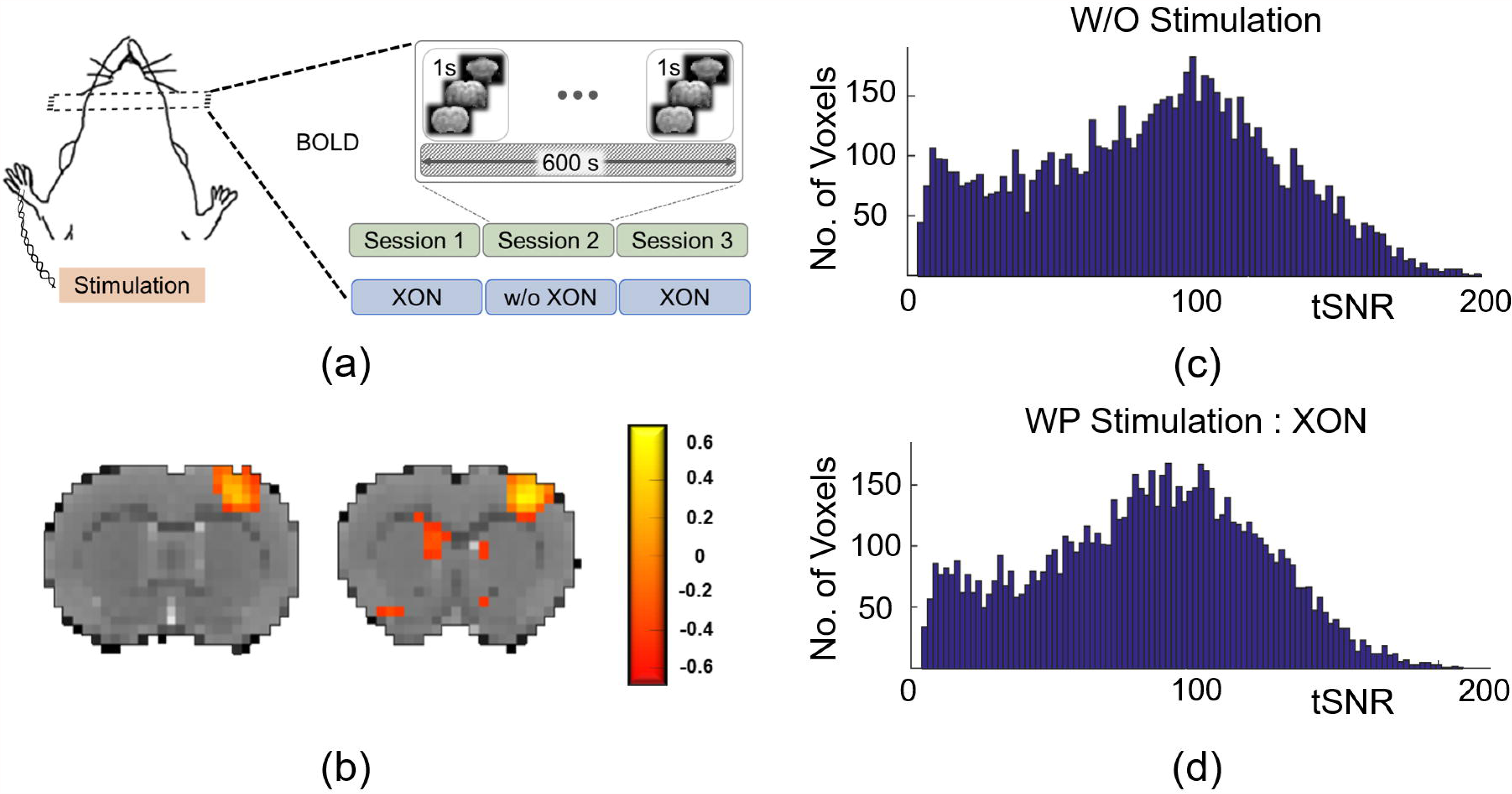
BOLD fMRI and Muscle Stimulation: (a) Simultaneous BOLD fMRI imaging paradigm alongside forepaw stimulation in a rat model. (b) Evoked BOLD response due to electrical stimulation. tSNR histograms based on fMRI data: (c) without XON stimulator inside MRI and (d) with XON stimulator providing stimulation while placed inside MRI bore.

### C. Gastric MRI and vagal nerve stimulation

Vagus nerve stimulation (VNS) has shown to influence gastric motility [40], [43], [44]. Fig. 7d shows one such modulatory effect of VNS, through XON stimulator, on stomach contraction. Stimulation of the nerve fibers were directed towards the stomach (efferent) through a polarity selective stimulation paradigm, shown in Fig. 7a. The cross-sectional area of the distal antrum (lower part of the stomach, adjoining to duodenum) was quantified from the acquired images through the procedure described in an earlier study [45].

**Fig. 6.**
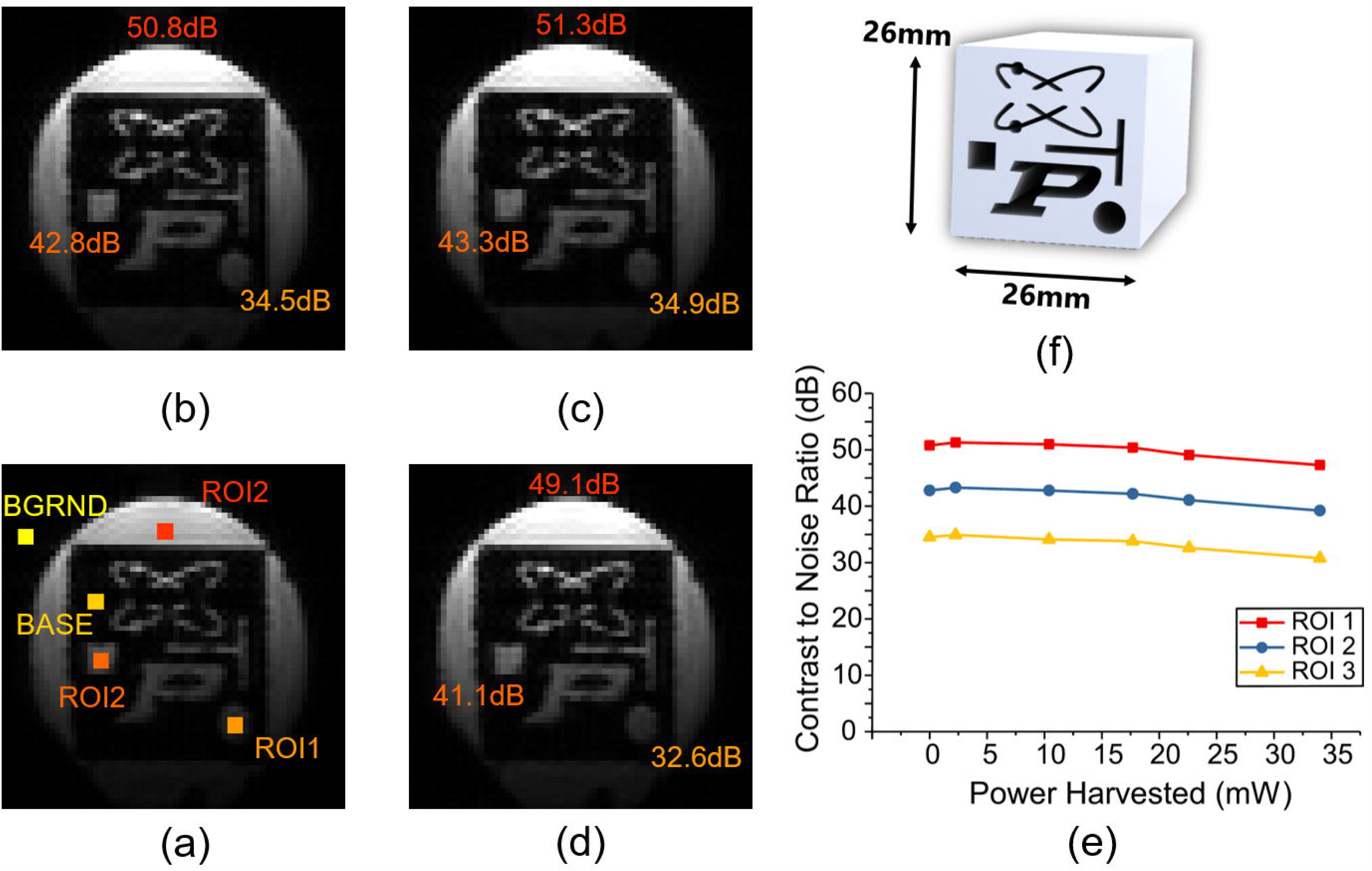
Gastric MRI and VNS: (a) Vagal nerve stimulation during concurrent gastric MRI. (d) Variation of antral cross-sectional area of the stomach in response to stimulation. MRI images acquired through FLASH scans of rat stomach (d) with no device operation and (c) while wireless power was har-vested by XON. CNR of images at region (x) for both images were calculated with respect to (y) [20log (μ_X_ -μ_Y_ /σ_B_)]

**Fig. 7.**
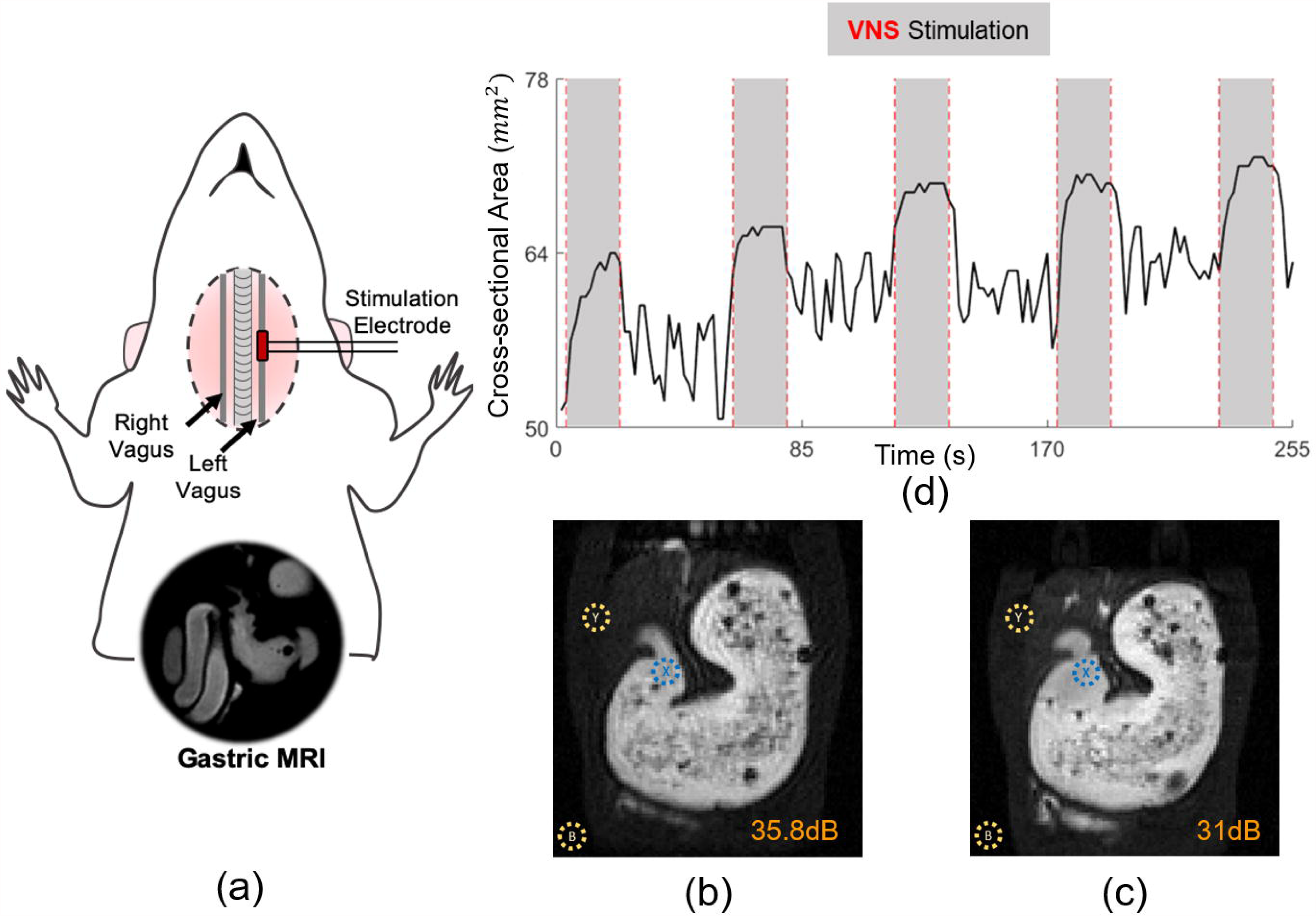
Effects on MRI data quality: Phantom images taken through EPI scans at harvested power levels of (b) 0mW, (c) 2.3mW and (d) 24mW. CNR of images computed at three ROIs (a) [20log (μ_ROI_ -μ_BASE_ / σ_BGRND_)] with respect to background. (f) 3D printed phantom model. (f) Variation of CNR at three ROIs with different harvested power levels.

The contraction amplitude in the distal antrum decreased significantly during stimulation zones as marked in the Fig. 7c. The 20S-ON-40S-OFF stimulation paradigm induced gastric secretion while damping contraction, which resulted in increased antral volume. This can be clearly identified as the average cross-sectional area increased after onset of each VNS. The inhibitory effect and following rebound contractions were reproduced over multiple cycles. The efferent VNS induced secretion is in agreement with previous findings [46], [47].

### D. Effects on MRI data quality

We tested the effects of device operation on the temporal signal-to-noise ratio (tSNR) of fMRI. tSNR elucidate the data quality of an acquired fMRI time series [48], [49] and was used to understand the effects of external elements introduced into the MRI (XON stimulator). Initially the tSNR for every voxel was calculated and plotted as a histogram as shown in Fig. 5. Figure 5c shows the tSNR observed when there was no device present in the MRI bore. Figure 5d shows the tSNR values when the device was providing simulation while harvesting power from the gradients.

CNR is another important parameter to quantify image quality as it provides knowledge regarding the ease of obtaining experimentally induced activation or contrast during imaging [49]. We obtained CNR of the EPI images during different levels of power harvesting through the X-ON device. As a surface coil was used to record these images, CNR was calculated at three regions of interest (ROI) to account for variation of coil sensitivity. At different levels of harvested power, the CNR of the images remained unaltered as shown in Fig.6. At higher power levels, slight decrease in CNR can be associated with increased image background noise. Additionally, no geometric distortion in the custom phantom (Fig. 6a) images was observed due to gradient power harvesting (Fig.6b-e).

CNR was also calculated for the FLASH images and were compared. Figure 7c shows the CNR at specific ROI while simultaneously the wireless stimulator scavenged energy from the gradients. Fig. 7b shows the CNR of same ROI when the device was not operating.

## Discussions

In this paper, we have presented a miniaturized stimulation system (XON), which is tailored for simultaneous operation along-side MR imaging. The unique plug-and-play system wirelessly synchronizes with the imaging system and does not require any additional cable connection. The small footprint of the system provides a safer solution for simultaneous stimulation and MRI at high magnetic fields. The power harvesting module described herein, can scavenge up to 72 mW power from the varying gradients present during typical MRI scans, e.g. fMRI.

### A. Design considerations

High voltage compliance during bipolar stimulation is essential during different tissue stimulation (i.e. muscle stimulation). Utilizing the H-bridge, the stimulator front end generated positive and negative load currents from a single harvested DC voltage. Thus, peak-to-peak load voltage during biphasic stimulation cycle may range larger than the actual harvested single supply DC value (+11 V). For this design architecture, the load voltage compliance can be expressed as

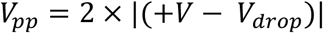

Considering +V = 11 V and V_drop_= 1 V peak-to-peak voltage compliance is 20 V, slightly less than twice the harvested supply voltage. The stimulator provided charge balanced current stimulation of up-to 20mA in the current setup. A peak instantaneous power delivery of 400mW can be achieved through the system, which is useful for large muscle or transcutaneous stimulation [50].

The presented stimulator can generate constant anodic-first, cathodic-first, symmetric and non-symmetric biphasic stimulation pulses. The device provides an optimal shape of the stimulation current pulse for necessary charge injection and its balancing. Maintenance of a good charge balance can be affected by large electrode voltage swings. To address this challenge, the presented architecture utilizes a linear mode circuitry and the same set of hardware components in the stimulator front-end for both positive and negative load currents. As a result, the absolute current magnitude repeats from first stimulation phase to the next phase and minimizes any mismatch error. Hence, accuracy of net charge injection remained unaltered by large voltage swings and long-term temperature variability.

Individual stimulation profiles are generated by the high voltage analog switches (Fig. 2: S1, S2 and S4). These switches are controlled by the onboard microcontroller. The stimulator can be operated in either MRI synchronized stimulation mode or a fixed frequency stimulation mode. Thus, the device can accommodate varied no of event driven or block driven experimental paradigms. For, the animal experiments we used two different block designs (a) 30s-ON-30s-OFF and (b) 20s-ON-40s-OFF with monophasic stimulation as shown in Fig. 4. Additionally, the MRI synchronized stimulation mode was also tested through phantom imaging as shown in Fig. 3e.

The same architecture can be used to generate bi-phasic stimulation pulses and insert inter-phase delays (IPD) without any hardware modification. The maximum stimulation frequency depends on the dynamics of the control signals, the timing characteristics of the stimulator front-end, and its output capacitance. The analog-front end of the stimulator was tested to operate between the frequency range of 0.1 Hz - 1000 Hz which is suitable for a wide variety of muscle and neuro-stimulation applications.

The base station provides a simple solution for stimulation studies, where stimulation blocks are independent to imaging triggers. During the such studies, MRI imaging triggers and stimulation blocks can be recorded on the same platform through the custom GUI. The sampling bandwidth is sufficient to accommodate these triggers reliably and accurately. These trigger information can be used later to correlate the stimulation and image trains and observe the effects of neuromodulation through the MRI images.

### B. Wireless power harvesting

The power harvesting coil was chosen to minimize any interference with the MRI receiver coils. The self-resonance frequency of the coil (134 kHz) was a few orders lower than the tuned frequency of the surface coil and the volume coil (~300 MHz). In addition, the small size (2 cm diameter) and separation from the receiver coils (6 cm) ensured a very low coupling coefficient. As a result, the image SNR remained unchanged even when relatively higher power (72 mW) was harvested by the system. Thus, the system did not require any dedicated high voltage RF switches to isolate itself from the MRI receiver/ transmission coil [51].

With the device harvesting energy from the MRI electromagnetic fields, it was imperative to understand quantitatively, the effect of energy harvesting on MRI image quality. The currents generated in the coil can be classified as eddy currents, which can deteriorate image quality with N/2 ghosting artifacts. However, analyzing different sequences (EPI and FLASH) and MRI images ghosting effect or geometric distortion due to such eddy currents were not observed. Additionally, no degradation of image quality was noticed while the tSNR and CNR of MRI images were compared between two conditions (a) while the device was harvesting power to operate and (2) while the device remained inoperative, outside the MRI-bore.

### C. Future development and application

In this study, we concentrated on proof of concept and possible applications of the novel MRI powered stimulator. The safety and compatibility of introducing such integrative stimulation modules remains to be fully investigated for future preclinical and clinical applications.

The low power design of the stimulator front-end allows for increase of stimulation channels without significant increase in the power budget (<1mw/channel). Thus, multi-channel stimulation capabilities can be implemented through the XON stimulator system in the future to enable simultaneous stimulation at multiple locations. Additionally, the large current delivery capabilities of the stimulator are well suited for additional modes of stimulation like optical [52] or mechanical stimulation [53], [54] during concurrent imaging studies. Another interesting approach may be to integrate a low power recording system [55], [56] with the power harvesting and stimulation unit described here. These integrative systems can enable simultaneous recording of biopotentials, electrical stimulation and imaging (MRI and fMRI) for closed loop image guided neuromodulation.

The miniaturized system was designed to be placed and operated within a 7 T animal MR-scanner. However, with a small foot print and relatively low-cost design the XON stimulator can be easily tuned to fit other animal MRI systems. It is well suited for basic-science and preclinical studies involving simultaneous imaging and neuromodulation. Future development of this technique is expected to enable MRI-based multimodal imaging for different neuroscience applications in both animal and human studies.

## Acknowledgement

This work was supported by funding from the National Institutes of Health (OD023847, NS105298, MH104402), Purdue University, MR-Link LLC, and Purdue Research Foundation.

## References

1. M. Kogan, M. McGuire, and J. Riley, “Deep Brain Stimulation for Parkinson Disease,” Neurosurgery Clinics of North America. 2019.

2. Dormont, D. Seidenwurm, D. Galanaud, P. Cornu, J. Yelnik, and E. Bardinet, “Neuroimaging and deep brain stimulation.,” Am. J. Neuroradiol., vol. 31, no. 1, pp. 15–23, 2010.

3. T. Taira, “Deep brain stimulation for dystonia,” in Deep Brain Stimulation for Neurological Disorders: Theoretical Background and Clinical Application, 2015.

4. B. M. Uthman et al., “Effectiveness of vagus nerve stimulation in epilepsy patients: A 12-year observation,” Neurology. 2004.

5. M. K. Lyons, “Deep brain stimulation: current and future clinical applications.,” Mayo Clin. Proc., vol. 86, no. 7, pp. 662–72, 2011.

6. M. Raslan, “Deep brain stimulation for chronic pain: Can it help?” Pain, 2006.

7. R. G. Bittar et al., “Deep brain stimulation for pain relief: A meta-analysis,” Journal of Clinical Neuroscience. 2005.

8. W. S. Gibson et al., “Functional correlates of the therapeutic and adverse effects evoked by thalamic stimulation for essential tremor.,” Brain, vol. 27, no. 40, pp. 10659–10673, 2016.

9. J. Cao et al., “Gastric stimulation drives fast BOLD responses of neural origin,” Neuroimage, 2019.

10. W. S. Gibson et al., “The Impact of Mirth-Inducing Ventral Striatal Deep Brain Stimulation on Functional and Effective Connectivity,” Cereb. Cortex, p. bhw074, 2016.

11. S. Dietrich et al., “A novel transcutaneous vagus nerve stimulation leads to brainstem and cerebral activations measured by functional MRI,” Biomed. Tech., 2008.

12. J. T. Narayanan, R. Watts, N. Haddad, D. R. Labar, P. M. Li, and C. G. Filippi, “Cerebral activation during vagus nerve stimulation: A functional MR study,” Epilepsia, 2002.

13. G. Schaefers, “Testing MR safety and compatibility,” IEEE Eng. Med. Biol. Mag., vol. 27, no. 3, pp. 23–27, 2008.

14. J. S. Thornton, “Technical challenges and safety of magnetic resonance imaging with in situ neuromodulation from spine to brain,” European Journal of Paediatric Neurology. 2017.

15. Kanal et al., “American College of Radiology White Paper on MR Safety,” Am. J. Roentgenol., vol. 178, no. 6, pp. 1335–1347, Jun. 2002.

16. Schaefers and A. Melzer, “Testing methods for MR safety and compatibility of medical devices,” Minimally Invasive Therapy and Allied Technologies. 2006.

17. Hovet, H. Ren, S. Xu, B. Wood, J. Tokuda, and Z. T. H. Tse, “MRI-powered biomedical devices,” Minimally Invasive Therapy and Allied Technologies. 2018.

18. E. K. Keeler et al., “Accessory equipment considerations with respect to MRI compatibility,” Journal of Magnetic Resonance Imaging. 1998.

19. B. A. Hargreaves, P. W. Worters, K. B. Pauly, J. M. Pauly, K. M. Koch, and G. E. Gold, “Metal-induced artifacts in MRI,” Am. J. Roentgenol., 2011.

20. Akbari, C. P. Rockel, D. A. Kumbhare, and M. D. Noseworthy, “Safe MRI-Compatible electrical muscle stimulation (EMS) system,” J. Magn. Reson. Imaging, 2016.

21. P. R. Kileny, T. A. Zwolan, S. Zimmerman Phillips, and S. A. Telian, “Electrically Evoked Auditory Brain-Stem Response in Pediatric Patients with Cochlear Implants,” Arch. Otolaryngol. Neck Surg., 1994.

22. Alwatban, C. N. Ludman, S. M. Mason, G. M. O’Donoghue, A. M. Peters, and P. G. Morris, “A method for the direct electrical stimulation of the auditory system in deaf subjects: A functional magnetic resonance imaging study,” J. Magn. Reson. Imaging, 2002.

23. C. Silva, S. P. Lee, G. Yang, C. Ladecola, and S. G. Kim, “Simultaneous blood oxygenation level-dependent and cerebral blood flow functional magnetic resonance imaging during forepaw stimulation in the rat,” J. Cereb. Blood Flow Metab., 1999.

24. D. L. Hayes, D. R. Holmes, and J. E. Gray, “Effect of 1.5 tesla nuclear magnetic resonance imaging scanner on implanted permanent pacemakers,” J. Am. Coll. Cardiol., vol. 10, no. 4, pp. 782–786, Oct. 1987.

25. J. Yeung, P. Karmarkar, and E. R. McVeigh, “Minimizing RF heating of conducting wires in MRI,” Magn. Reson. Med., 2007.

26. M. Meinzer, R. Lindenberg, R. Darkow, L. Ulm, D. Copland, and A. Flöel, “Transcranial Direct Current Stimulation and Simultaneous Functional Magnetic Resonance Imaging,” J. Vis. Exp., 2014.

27. Y. Cabral-Calderin, K. A. Williams, A. Opitz, P. Dechent, and M. Wilke, “Transcranial alternating current stimulation modulates spontaneous low frequency fluctuations as measured with fMRI,” Neuroimage, 2016.

28. R. G. Garcia et al., “Modulation of brainstem activity and connectivity by respiratory-gated auricular vagal afferent nerve stimulation in migraine patients,” Pain, 2017.

29. K. A. Williams, Y. Cabral-Calderin, C. Schmidt-Samoa, C. A. Weinrich, P. Dechent, and M. Wilke, “Simultaneous Transcranial Alternating Current Stimulation and Functional Magnetic Resonance Imaging,” J. Vis. Exp., 2017.

30. S. Assecondi, C. Lavallee, P. Ferrari, and J. Jovicich, “Length matters: Improved high field EEG-fMRI recordings using shorter EEG cables,” J. Neurosci. Methods, vol. 269, pp. 74–87, 2016.

31. T. L. Yearwood and L. T. Perryman, “Peripheral neurostimulation with a microsize wireless stimulator,” Prog. Neurol. Surg., 2015.

32. J. Lee, J. Jang, and Y. K. Song, “A review on wireless powering schemes for implantable microsystems in neural engineering applications,” Biomedical Engineering Letters. 2016.

33. W. Turner, C. M. Tang, B. Beck, T. H. Mareci, and R. Bashirullah, “Nuclear magnetic resonance energy harvesting for ultra-low power biomedical implants,” in 2011 IEEE 12th Annual Wireless and Microwave Technology Conference, WAMICON 2011, 2011.

34. J. Höfflin, E. Fischer, J. Hennig, and J. G. Korvink, “Energy Harvesting with a figure-8 coil - towards energy autonomous MRI detection,” no. April 2013, 2016.

35. L. G. Hanson, T. E. Lund, and C. G. Hanson, “Encoding of electrophysiology and other signals in MR images,” J. Magn. Reson. Imaging, vol. 25, no. 5, pp. 1059–1066, 2007.

36. Weissler et al., “PET/MR synchronization by detection of switching gradients,” IEEE Trans. Nucl. Sci., vol. 62, no. 3, pp. 650–657, 2015.

37. R. W. Cox, “AFNI: Software for analysis and visualization of functional magnetic resonance neuroimages,” Comput. Biomed. Res., 1996.

38. M. Jenkinson, C. F. Beckmann, T. E. J. Behrens, M. W. Woolrich, and S. M. Smith, “FSL - Review,” Neuroimage, 2012.

39. J. Cao, K. H. Lu, T. L. Powley, and Z. Liu, “Vagal nerve stimulation triggers widespread responses and alters large-scale functional connectivity in the rat brain,” PLoS One, 2017.

40. K. H. Lu et al., “Vagus nerve stimulation promotes gastric emptying by increasing pyloric opening measured with magnetic resonance imaging,” Neurogastroenterol. Motil., 2018.

41. K.-H. Lu, J. Cao, R. Phillips, T. L. Powley, Z. Liu, and P. Research Associate, “Differential Effects of Afferent and Efferent Vagus Nerve Stimulation on Gastric Motility Assessed with Magnetic Resonance Imaging.”

42. J. E. B. Randles, “Kinetics of rapid electrode reactions,” Faraday Discuss., 1947.

43. B. E. Lacy and K. Weiser, “Gastric motility, gastroparesis, and gastric stimulation,” Surgical Clinics of North America. 2005.

44. E. Soffer, “Gastric electrical stimulation for gastroparesis,” Journal of Neurogastroenterology and Motility. 2012.

45. K.-H. Lu, J. Cao, S. Oleson, M. Ward, T. Powley, and Z. Liu, “Vagus Nerve Stimulation Promotes Gastric Emptying in Rat Measured by Magnetic Resonance Imaging,” in Proc. Joint Annual Meeting ISMRM-ESMRMB, Paris, France, 2018.

46. Grundy and T. Scratcherd, “Effect of stimulation of the vagus nerve in bursts on gastric acid secretion and motility in the anaesthetized ferret.,” J. Physiol., 1982.

47. R. Berthoud, W. B. Laughton, and T. L. Powley, “Vagal stimulation-induced gastric acid secretion in the anesthetized rat,” J. Auton. Nerv. Syst., 1986.

48. C. Triantafyllou et al., “Comparison of physiological noise at 1.5 T, 3 T and 7 T and optimization of fMRI acquisition parameters,” Neuroimage, 2005.

49. M. Welvaert and Y. Rosseel, “On the definition of signal-to-noise ratio and contrast-to-noise ratio for fMRI data,” PLoS One, 2013.

50. D. Karpul, G. K. Cohen, G. D. Gargiulo, A. van Schaik, S. McIntyre, and P. P. Breen, “Low-power transcutaneous current stimulator for wearable applications.,” Biomed. Eng. Online, vol. 16, no. 1, p. 118, Oct. 2017.

51. K. Byron, F. Robb, P. Stang, S. Vasanawala, J. Pauly, and G. Scott, “An RF-gated wireless power transfer system for wireless MRI receive arrays,” Concepts Magn. Reson. Part B Magn. Reson. Eng., 2017.

52. P. Lin, Z. Fang, J. Liu, and J. H. Lee, “Optogenetic Functional MRI,” J. Vis. Exp., 2016.

53. L. Li, C. Weiss, A. C. Talk, J. F. Disterhoft, and A. M. Wyrwicz, “A MRI-compatible system for whisker stimulation,” J. Neurosci. Methods, 2012.

54. Lu et al., “Low-but Not High-Frequency LFP Correlates with Spontaneous BOLD Fluctuations in Rat Whisker Barrel Cortex,” Cereb. Cortex, 2016.

55. R. Mandal, N. Babaria, J. Cao, and Z. Liu, “Adaptive and Wireless Recordings of Electrophysiological Signals During Concurrent Magnetic Resonance Imaging,” IEEE Trans. Biomed. Eng., vol. 66, no. 6, pp. 1649–1657, Jun. 2019.

56. R. Mandal, N. Babaria, J. Cao, and Z. Liu, “Recording Electrophysiological Signals Through MR Receiver Coils During Concurrent fMRI,” Proc. Intl. Soc. Mag. Reson. Med. 26 4660, 2018.

